# Neuron particles capture network topology and behavior from single units

**DOI:** 10.1101/2021.12.03.471160

**Authors:** Gaurav Gupta, Justin Rhodes, Roozbeh Kiani, Paul Bogdan

## Abstract

While networks of neurons, glia and vascular systems enable and support brain functions, to date, mathematical tools to decode network dynamics and structure from very scarce and partially observed neuronal spiking behavior remain underdeveloped. Large neuronal networks contribute to the intrinsic neuron transfer function and observed neuronal spike trains encoding complex causal information processing, yet how this emerging causal fractal memory in the spike trains relates to the network topology is not fully understood. Towards this end, we propose a novel statistical physics inspired neuron particle model that captures the causal information flow and processing features of neuronal spiking activity. Relying on synthetic comprehensive simulations and real-world neuronal spiking activity analysis, the proposed fractional order operators governing the neuronal spiking dynamics provide insights into the memory and scale of the spike trains as well as information about the topological properties of the underlying neuronal networks. Lastly, the proposed model exhibits superior predictions of animal behavior during multiple cognitive tasks.

## Main

One of the general principles of most complex systems is that they consist of numerous structures or units with dynamic inter-dependence and interactions among them. The complex network interactions can yield unique output patterns, that can result in higher-order functional responses. For example, neural population interactions leading to thoughts and movements (*1*), social interactions in cyber-terrorist networks (*2*), ecological interactions between species (*3*), interactions between climate features (*4*). In practical settings, we only have partial knowledge of the structure, e.g., knowledge of only a fraction of the nodes in the complex network. Herein, we address the question of inferring the underlying network topology using events recorded from multiple single nodes that make up only a small fraction of the total nodes in the network. In this pursuit, the network of neurons is a prominent example which is the primary case study in the current work.

The human brain functions through networks of interactions between billions of neurons, glia and vascular systems connected through a complex architecture (*5–7*). Together, these systems enable shifts in electrical activation patterns across brain networks, depending on sensory stimulation and intrinsic states which are shaped by past experience (*8*). The view that all our perceptions, thoughts, emotions, and goal directed behaviors can ultimately be traced to the dynamic changes in the network activity within the brain is widely held (*5, 8–13*). However, to date, no method is capable of measuring all the relevant features to decode the network dynamics into units of behavior. While technologies that analyze anatomical connections and proxies of neuronal activity across the entire brain such as magnetic resonance imaging (MRI) and functional MRI (fMRI) can be used to infer certain aspects of macroscopic (brain region) network dynamics, information is lost during the summation (fusion) that could be crucial, such as timing of intervals between spikes (*12, 14, 15*). While technologies for measuring action potentials or spikes from single neurons has advanced considerably in recent years (*16, 17*), even the best technologies can only measure hundreds to thousands of units usually only in a few restricted regions of the brain(*18*) and providing very scarce information about the neuronal networks involved in the respective cognitive functions.

Hence, there is a need for developing new mathematical methods capable of inferring features of the neuronal network from measuring only a small sample of single units (from a few to thousands) and relate the statistical characteristics of neuronal spiking activity with the overall brain functionality and behavior.

One of the major limitations of current statistical models of neuron spike data is that they do not account for the heterogeneity and non-Markovianity in the spike trains from individual units. We hypothesize that neuronal spike trains could be modeled more precisely by fractional order partial differential equations. Further, we hypothesize that a more precise statistical representation of the spiking patterns that captures the fractal properties and long-range memory of the inter-spike intervals could uncover features about the underlying network topology and dynamics, and thereby provide new insight into how cognitive and sensory processes are represented in the brain. The need for non-Markovian fractal methods is recognized as an important open-problem in the neuroscience community (*19*). The current work, by proposing a novel fractional diffusion-based analogy for the neural spike trains, is a step towards addressing this challenge.

Currently, the modeling of neural computations can be categorized into first order integrate and fire (I&F) dynamic models (*20–24*)and renewal theory inspired refractory density models (*25, 26*)(RDMs). The I&F models work under the assumption of homogeneous neural populations and conceive the neuronal dynamics as a summation (hence, the integrate terminology) of nonlinearly filtered action potentials (above a critical voltage threshold). The I&F models consist of integer-order differential equations of the population activity that pass the homogeneous neuronal input potentials through a nonlinear filter. Although successful to model cortical visual processing (*27–29*), motor control and decision making(*30*), the I&F models cannot capture the heterogeneity, non-Markovianity and microscopic non-Gaussian statistics of neuronal dynamics. In contrast, the RDM models determine the evolution of a time dependent probability of observing a spike after a certain amount of time elapsed since the last spike through an integer order partial differential equation capturing the interdependence between the inter-spike interval (i.e., the time since the last spike) gradient and the neuronal population firing activity subject to a nonlocal boundary condition. However, the RDM models cannot account for the heterogeneity and non-Markovianity of neuronal populations or shed light on the type of network topology characterizing specific emerging neuronal circuits during cognitive tasks.

Unlike existing work that assumes that the neuronal spike trains are Markovian and models their activity through the Poisson point processes formalism, we show that the inter-spike intervals are more regular, possess long-range memory and fractal characteristics and their statistics are better described by fractional order partial differential equations. Moreover, we show that the microscopic statistics encode emergent macroscopic topological properties about the underlying network, and display superior predictions of animal behavior during multiple cognitive tasks. Through simulations, we further discover how the macroscopic neural network activity influences the microscopic statistics of neuronal spiking dynamics. This framework formalizes, generalizes and unifies the mathematical modeling of microscopic neuronal dynamics while opening new analytical avenues for cognitive and sensory computational neuroscience including detecting and studying cognitive and sensory abnormalities or psychiatric disorders.

### Neuron particles model

We introduce a novel concept of *neuron particles* to explain the causal fractal memory and scale component in the spike trains. We define the inter-spiking intervals (ISI) in the spike trains as neuron particles of the size corresponding to the length of ISI (Fig.1a). As shown in Fig.1b, the neuron particles start from the origin of space-time and make a jump when the associated ISI recur in the spike train at time *t*. The jump made by a particle in space is a fixed proportion of its size. Therefore, smaller particles make smaller jumps and bigger particles bigger jumps. The procedure to obtain the neuron particles, Neuron Particles Process (NPP), is outlined in Fig.2 (for details see Methods). The neuron particles traverse trajectories *X*(*t*), which we model as realization of a multi-parameter fractional partial differential equation (*31, 32*)(FPDE) for each spike train (see Fig.1c). The space-time fractional diffusion equation captures the scale and fractal memory using fractional Riesz-Feller (*33*)(order *α*, skewness *θ*) and Caputo time-fractional (*34*)(order *β*) derivative, respectively. The trajectories are used to estimate the parameters of the FPDE using the fractional moments algorithm (*31*).

**Fig. 1.**
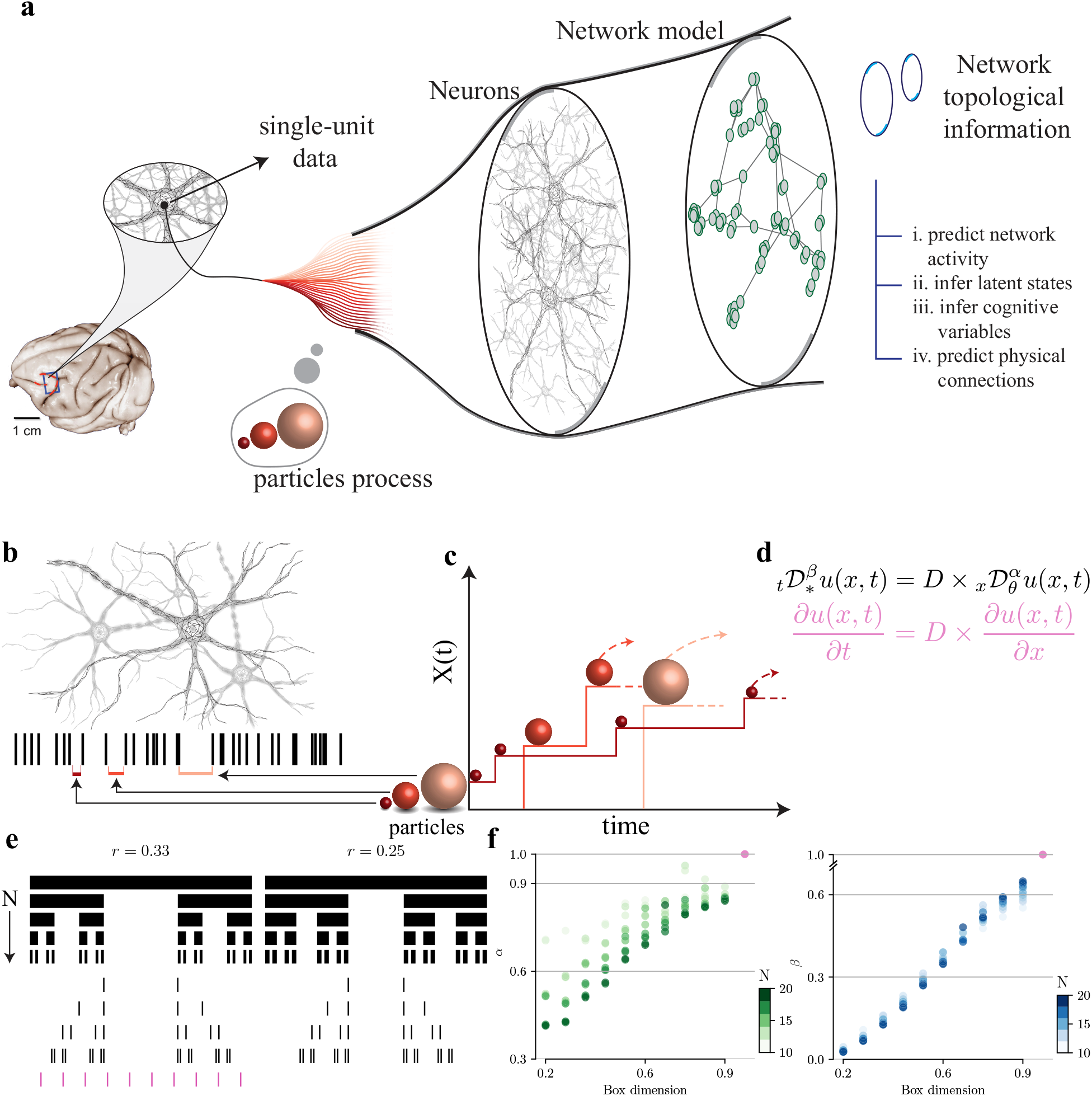
Neuron “particles” capture the statistical complexity of neuronal spike trains. a, The particles process aims at extracting the high-dimensional information from a single-unit events data. The high-dimensional information is used to identify the network topological information. Additionally, the particles process can be used to perform the tasks as mentioned in (i)-(iv). In the present work, the task (iii) is discussed as Monkey’s choice prediction. b, Neuron action potentials (spike trains) are recorded from a single unit. The inter-spiking intervals (ISI) are identified as “particles,” illustrated by spherical symbols of various sizes. Image credit: shutterstock.com. The ISI are divided into discrete bins as explained in the Methods. c, Each particle starts at the origin and stays stationary until it recurs in the neural spike train, after which it makes a jump proportional to the length of ISI resulting in trajectory *X*(*t*). d, The space-time particle trajectories are modeled through a multi-parameter fractional partial differential equation (PDE) which captures the scale and memory through *α* and *β* parameters, respectively. Since *X*(*t*) ≥ 0, the skewness parameter *θ* = −*α* in neuron particles setup. e, Cantor set example to generate scale-free spike train patterns. The deletion ratio (*r*) can vary from 0 to 1 (0<r<1) and division factor *N* > 1. A special case of uniform spike trains with equal ISI across time in pink color. f, Estimated fractional PDE parameters (*α, β*) for Cantor spike trains with different box dimensions. The box dimension is the slope of the logarithm of the number of ISI of size *ϵ*, i.e., log (*N*(*ϵ*)) and the logarithm of the inverse ISI size, i.e., log(1/*ϵ*). For a Cantor-like spike train, resulting from the corresponding Cantor set, the box dimension is −log(2)/log((1-r)/2). The special case of uniform spikes in pink in d can be represented as a differential equation in pink in c which has *α* = *β* = 1. The estimated values of (*α, β*) for uniform spikes is pink dot in e.

**Fig. 2.**
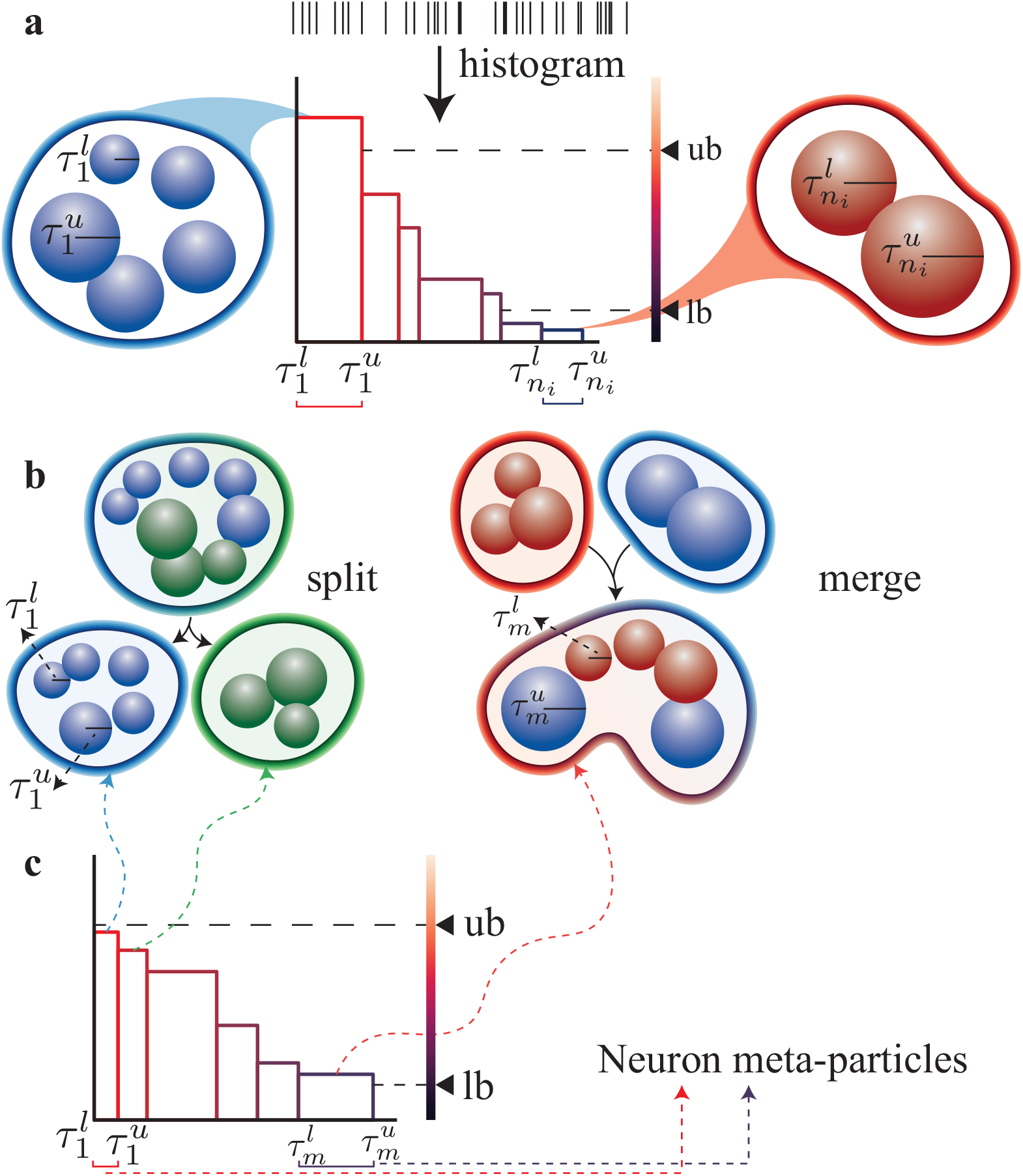
Neuron particles process. a, A histogram of the inter-spike intervals (ISIs) from a single unit spiking data (see the spike train on top) is generated. Each interval is treated as a group of particles with the minimum radius 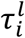 and the maximum radius 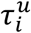. Two specified parameters ‘ub’ and ‘lb’ determine the upper- and lower-bound on the number of particles in each group. b, Groups with a larger number of particles than the upper bound are broken into smaller groups, while smaller groups are combined using ‘split’ and ‘merge’ steps, respectively. The iteration continues until each particles’ group has a number of particles in the range [lb, ub]. c, In the resulting histogram of ISIs, the ISI intervals are termed ‘neuron meta-particles’ and are such that the count of particles within them are in the range [lb, ub]. In the current work, we refer to the “neuron meta-particles” simply as “neuron particles”.

The multiscale phenomena and causal fractal memory captured by neuron particles in a spike train can be understood and validated by the hypothetical concept of Cantor spikes. Cantor sets (*35*) are formed by repeated deletion of a fixed ratio *r* from a unit interval [0,1] as shown in Fig.1d. The Cantor sets have numerous remarkable properties, and particularly useful in our case, is its fractal nature (statistical self-similarity). We generate Cantor spikes such that the successive ISIs are deleted segments of a Cantor set from left to right for given depth *N* and ratio *r*. The estimated values of scale and fractal memory parameters using neuron particles concept for Cantor spikes are shown in Fig.1e. The box dimension captures the fractality of Cantor spikes. We note that different depth *N* generate different lengths of Cantor spikes (2^*N*^) as well as different neuron particles (*r, rρ,* …, *rρ*^*N* − 1^), where *ρ*=(1-*r*)/2. From Fig.1e, we see that the fractal memory parameter *β* primarily is a variant of the fractality measure (box dimension) while estimations for various depths *N* cluster together. Therefore, the memory *β* captures fractal repeating patterns at various scales and does not depend on the depth. The overlap in scale parameter *α* of Cantor spikes across different box dimensions by varying *N* is due to common neuron particles among different (*r, N*) pair. Finally, a special case of uniform spikes is shown at the bottom (in pink) of Fig.1d where the ISIs are equal. Since the increments, or ISIs, are constant across time, the uniform spike train is straightforwardly modeled as a first order linear differential equation as in Fig.1c. Also, by using the neuron particles concept, we see that for the uniform spikes, the estimated value of *α* = *β* = 1 in Fig.1e (pink dot) represents first order differentials for both space and time.

### Topological features of neuronal networks influence the microscopic neuronal dynamics

The complex patterns in the neuron spikes are the result of the individual neuron transfer function and the network feeding into each neuronal unit (*36, 37*). An individual neuron is part of multiple closed-loops of different lengths that induce a complex pattern of recurring ISIs. By capturing the multi-scale repeating patterns in the ISIs, from the network perspective, *β* is related to the cumulative effect of multiple closed-loops around a neuron in the network. The network has influence on the statistics of the spike train (*1, 38*). The scaling of path lengths from a neuron to its neighbors at different hops can be captured using node-based multifractal spectrum. The scale of ISI which is represented using *α* is intuitively related to the node-based multifractal dimension.

We studied the effects of a wide range of complex network topologies on the fractal memory and scaling properties of a single neuronal spike train. The variation of fractal memory parameter *β* with different network topologies is shown in Fig. 3. For Erdös-Rényi (*39*)(ER) model of a random network (see Methods), we see in Fig. 3a that as the network becomes denser from the sparse state (*p* = 0.05) the memory parameter *β* of the ensemble increases and then decreases. A saturation occurs for dense networks with *p* > 0.2. The large number of connections provides a large input current to the neurons, thus, saturating the spiking patterns. The existence of a maximum point in the memory parameter shows that for some topologies (near *p* = 0.08), the network spikes are in a structured fashion. Next, in Fig. 3b, we see that for scale-free Bárabasi-Albert (BA) networks, the fractal memory decreases as outward links (*m*), independent of total number of neurons in the network, increases. Finally, for multi-fractal network (MFN) in Fig. 3c, the trend of fractal memory is inversely proportional to length measure *l* for depth *K*=2. However, as depth *K* increases the network gets sparser, and for *K*=4 the fractal memory attains a peak for *l* near 0.5 and then decreases. Taken together these results suggest that critical information about the type of network feeding into the single unit spike trains can be deduced from analyzing the spike trains using the neuron particle method. The information is not complete, but over certain values of *α, β*, and assuming a finite range of networks, we can describe the probability of features of the underlying network given a single spike train. To the best of our knowledge this has never been accomplished before. Such a probabilistic representation of the topology of the underlying network enables predictions of neural dynamics and behavior.

**Fig. 3.**
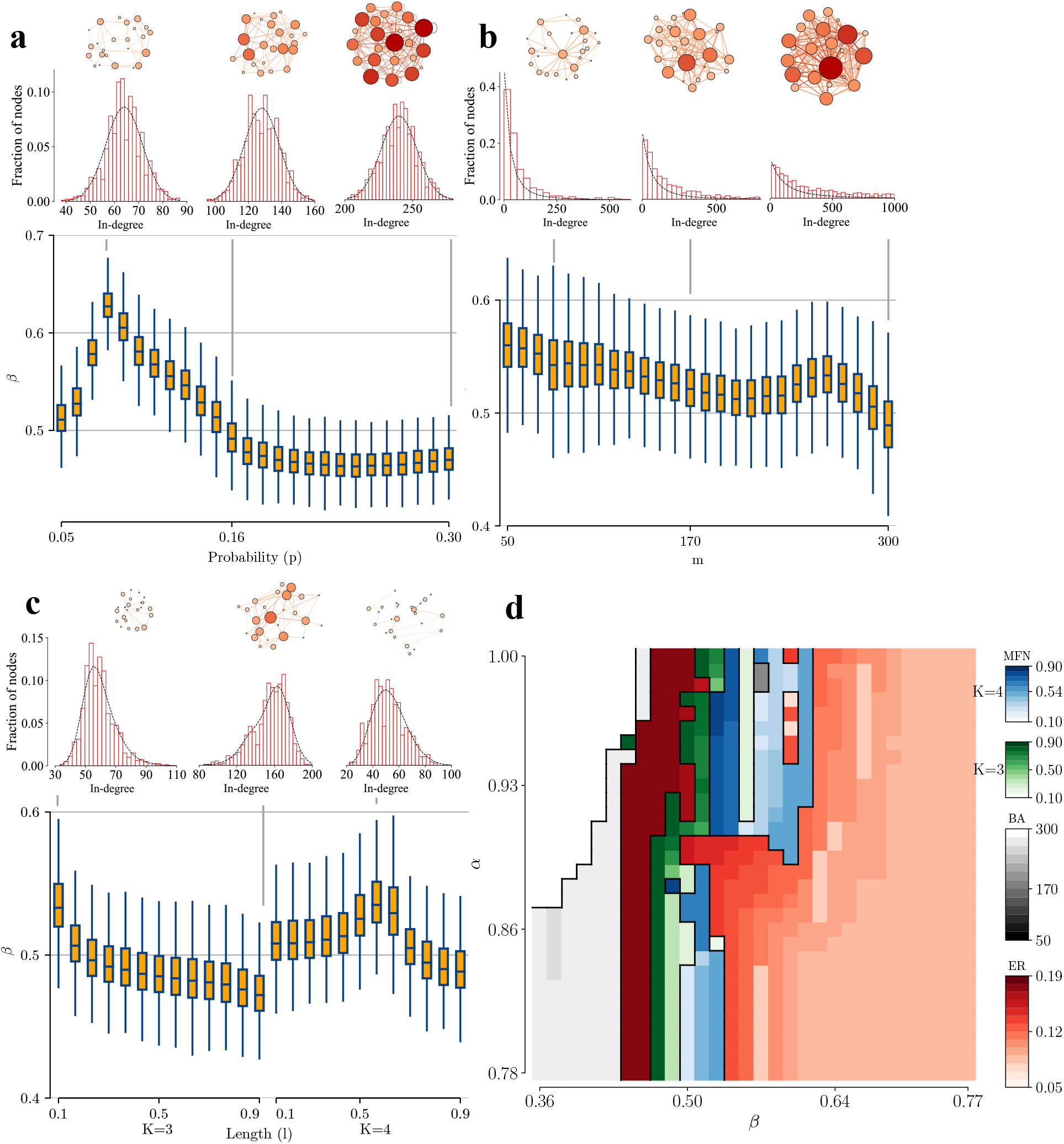
Network topologies influence the memory of neuronal dynamics. a, An Erdos-Renyi random network with probability of existence of edge being *p*. The probability varies linearly from 0.05 to 0.30 and consequently increases the average node-degree. The excitatory neuron node-degree scaled histogram is in red bars and the black curve fits the histogram with Gaussian distribution. The variation of the excitatory neuron *β* with respect to *p* is represented as box plots. The line across the box plot indicates the median, and upper and lower hinges indicate 75^th^ and 25^th^ percentiles (making IQR), respectively. The whiskers extend to the lowest and the largest values but within 1.5xIQR. b, Dorogovtsev (*48*) extension of Bárabasi-Albert (*47*) scale-free network with outgoing links *m* and attraction parameter *a=m*. The excitatory neuron node-degree scaled histogram is shown in red bars and the black curve fits the histogram using analytical expression (*48*). The outgoing links *m* varies from 50 to 300, and the variation of excitatory population fractal memory parameter *β* is shown as box plots. c, Multi-fractal network (*49, 50*)with base measure *P* = [0.6,0.5;0.5,0.4] (*m*=2) and length measure *L* = [*l*, 1-*l*] with *l* variation linearly from 0.1 to 0.9 for two values of depth *K* = 3, 4. The excitatory neuron node-degree scaled histogram is in red bars and the black curve fits the histogram with the analytical expression of degree distribution (*49*). The excitatory neuron population fractal memory parameter *β* is represented as box plots. d, Prediction spectrum of the networks using *α* and *β* computed from a single spike train. For each tuple (*α, β*) the most likely network is indicated by its corresponding color. Details regarding confidence of each network prediction is shown in the Fig. S2. For each network type (topology as well as parameters), 50 random realizations of networks were generated, and the estimated parameters for the neuron populations were concatenated for box plots.

Using the observed variation in the fractal memory parameter *β* (Fig. 3a-c) and scale parameter *α* (Fig. S1a-c), a prediction spectrum is constructed in Fig. 3d (full version in Fig. S2b). For each constituent networks ER, BA, and MFN with various parameter ranges, we show the most likely network that an estimated tuple (*α, β*) belongs. In the low values of *α* and *β*, the most probable network is BA with *m*~300, and ER with high connection probability (~0.19). On the contrary, high values for both parameters indicate ER with low connection probability. For some tuple values, almost all networks are equiprobable, and hence, it is difficult to predict a single one.

Multifractal spectrum analysis (MFSA) (see Methods) of the neuronal networks with weighted synapses identifies the higher-order connectivity patterns by averaging how the network grows around each node. For example, MFSA for an unweighted straight-line network yields a delta function around 1 (mono-fractal). For a general weighted network, MFSA represents a spectrum of fractal dimensions. For the ER, BA, and MFN networks considered in Fig. 3 and the results in Fig. S4b,c,d, we observe that both the (*α, β*) parameters characterizing the dynamics on a networked architecture and the dominant fractal dimension (corresponding to this highest value in the multifractal spectrum, see Methods) of the network exhibit similar qualitative patterns with varying network model parameters (i.e., *p* for ER, *m* for BA and (*l, K*) for MFN). For example, the ER with *p*=0.08 serves as a change-point where the trend alters in both *β* and dominant fractal dimension. A similar monotonic trend (but with different signs) is observed for the BA networks, For MFN (*K*=3), a monotonic trend with different sign is observed, while for *K*=4, the *l*=0.6 alters the trend in both *β* and dominant fractal dimension. This empirical evidence shows that the proposed (*α, β*) parameters, which can be estimated from spike trains, provide information about the network topology on which neuronal dynamics unfolds.

### Neuron particles predict behavior

To determine the extent to which the neuron particle analysis of single unit data is capable of predicting the behavior of an animal during a cognitive task we analyzed a few different publicly available data sets. In the first data set, hundreds of single units were recorded simultaneously in the pre-arcuate gyrus of Macaque monkeys while they were performing a direction discrimination task (*40*). Briefly, on each trial, the monkey views moving dots on a screen for 800 ms. A fraction of dots moves coherently in the same direction towards one target or another (T1 or T2) while the rest of the dots are displaced randomly (Fig. 4a). After the lapse of a random delay period, upon receiving a Go cue, the monkey makes a saccadic eye movement towards the target T1 or T2, where it thinks the dots were moving. We compared the prediction accuracy of the monkey’s choices based on a commonly used mean firing rate approach versus the neuron particles approach (Fig. 4b). We found a performance gain, with an averaged difference of 5.75% around the Go cue. Around the saccade initiation time, the spikes are distinct enough across the units such that both the firing rate and neuron particles trajectories perfectly predict the choice. However, neuron particles consistently outperform during motion viewing and delay periods.

**Fig. 4.**
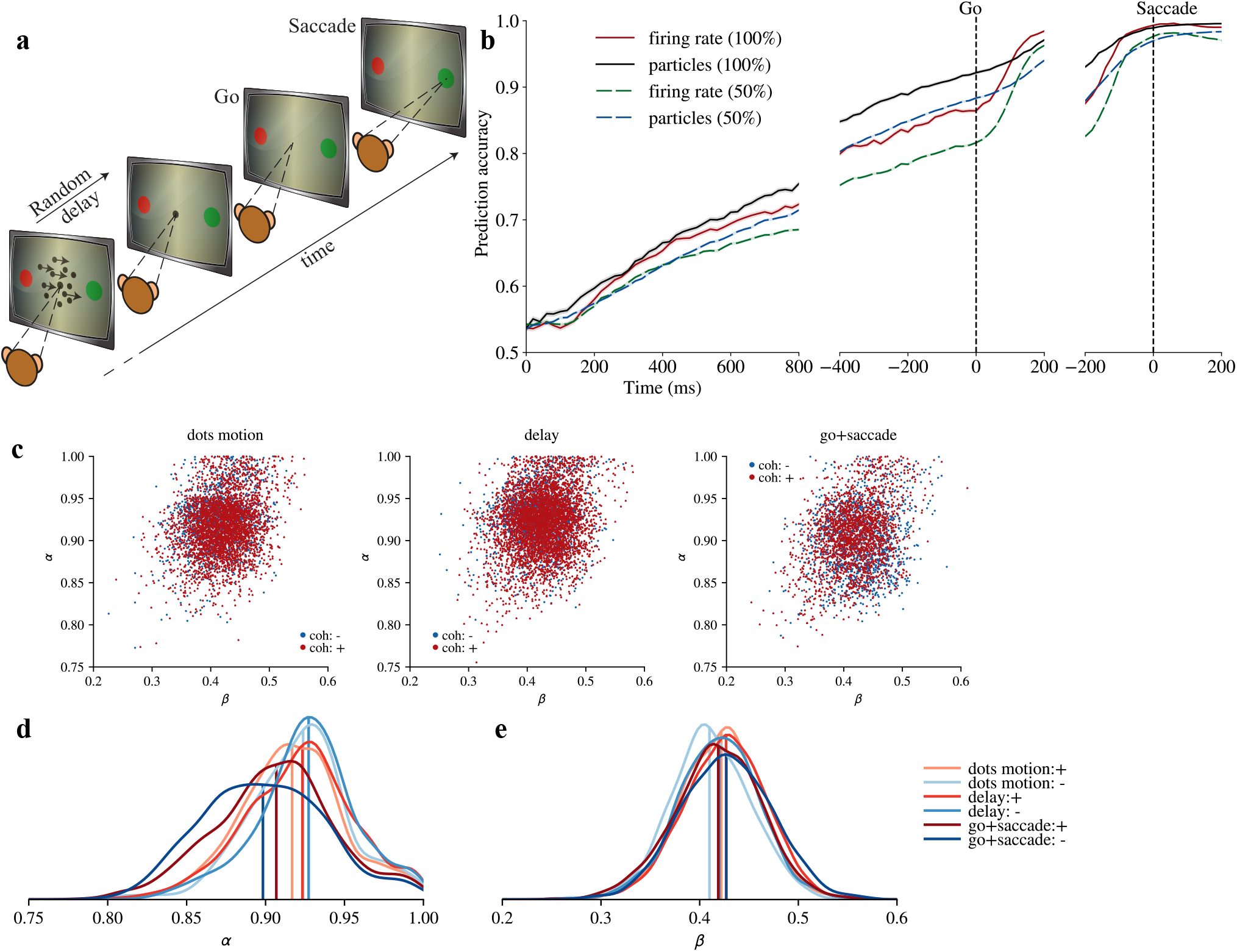
Neuron particles predict Monkey’s choices. a, Direction discrimination task by the animal. b, A comparison of choice prediction using recordings of neuron ensembles for neuron particles and mean firing rate approach. Both approaches use logistic regression with linear kernel and train/test split as 90/10. The particles trajectory is sampled every 20ms, and the firing rates are computed in a sliding 100ms bin with 20ms slide. The mean accuracy ± s.e.m. (across sessions and resampling) is indicated in dark and shaded colors. An additional case of 50% is considered, where to mimic the reduced data scenario, out of total, 50% uniformly random sampled units are taken, respectively. The sampling is done 5 times independently. c, The estimated parameters for monkey 1 across three epochs: motion viewing (200-800 ms after motion onset), delay (200 ms after dots offset to −50 ms from go cue), and go+saccade (go cue to 200 ms after saccade initiation). The parameter tuple was separately calculated for motion in favor and against the neuron’s preferred saccade (+ and – motion coherence, respectively). Similar results were obtained for monkey 2 (Fig. S8). d, e Gaussian-kernel smoothed densities for *α, β* across three epochs and 2 coherence levels with straight line being the median.

In an additional experiment, we randomly (uniform) sampled only 50% of the units to mimic a partial knowledge setup and predicted the monkey’s choice from the reduced data. The difference between firing rate and particles approach around the Go cue is 6.74%. With half the population, the prediction accuracy gap further increases. Another partial data setup is presented in the Fig. S7, where for each single unit we randomly removed some of the spiking event data. We see that when using only 50% of the data from each unit, the neuron particles approach suffers 1.76% performance loss, while mean firing rate suffers 4.70% loss at Go cue. Next, on using only 25% of the spiking data from each unit, the performance loss of neuron particles/mean firing rate is 4.52%/11.23% at Go cue. The fractality in the spike trains is less affected if some spikes are removed. The particles approach extracts more information from the same data which in turn can be utilized to better discriminate the spike patterns over time to predict the monkey’s choice.

### Observing a fraction of units in the network

Technological and practical challenges limit us to only partial observations of neuronal networks in most experimental setups, because often only a fraction of neurons in a circuit can be simultaneously recorded. Consequently, a natural question is whether the proposed (*α, β*)-fractional PDE analysis can provide robust information about the neuronal network topologies from partial and noisy neuronal dynamics observations (i.e., only a few spike trains). Towards addressing this question, we exploit the Bayesian graph model in Fig. S2a and investigate whether a given property of the estimated (*α, β*) pair provides information about the type of topology characterizing the neuronal network under varying degrees of network observability. With such property, only a partial sampling of the neurons in the network would enable accurate inference of the underlying network type. For a partial sample *δ*% of the total neurons in the network sampled randomly (uniform distribution), the relative deviation is defined as 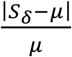, where *S*_*δ*_ is the empirical mean of the partial sample, and *µ* is the population mean. We compute the probability of relative deviation with various values of partial sample (*δ*) for networks in Fig.3 (Fig. S5, S6).

For ER networks with *p* = 0.08, by sampling only 1% of the neuronal population, the relative error in *α*/*β* is 3.09/5.63% with 90% confidence. For BA networks with *m*=300, the 1% sampling gives relative error of 5.96/10.26% with 90% confidence. The estimation analysis for BA networks with partial sampling has more deviation than ER and MFN models, a difference expected from the thick-tail of the connectivity distribution in BA networks. Taken together, these results suggest that a relatively small sample is needed to identify the underlying network with reasonable confidence. Moreover, these results show that the size of the sample depends to some extent on the network, with certain network structures requiring fewer units for the same identification confidence than others. Such information would be exceedingly useful for designing experiments where single unit data are collected as it can inform the minimum number of units that are needed to obtain a good representation of the underlying network structure.

### Scale and memory parameters are invariant to network size

The features used to infer the network topological structure from a single unit should ideally be independent of the network size. Additional experiments in Fig. S3 suggest that the scale (*α*) and fractal memory (*β*) parameters are indeed invariant to the network size under consideration. In Fig. 2, the network size is *N*=1000 and we take this as baseline to observe the network size effects upon changing *N*. By varying the network size from *N*=1000 to *N*=10000, the patterns in *α* and *β* are relatively stable for each network topology. This shows that the topology shapes the fractality of each network unit, independently of its size. The neuron particles, by capturing such fractality, can differentiate among different topological structures to a certain extent. We note that, for large BA networks with higher values of *m*, many connections occur for some nodes, and thus parameter estimates are variable due to massive incoming currents in the hub nodes. For the smaller networks, to maintain the same topological setup, the paucity of stimulating currents leads to a reduction in the spiking activities of the neurons. Therefore, for smaller networks the extracted network fractality patterns saturate and diverge from the bigger networks. The patterns quickly change for all three network types between *N*=250 to *N*=750. For ER and MFN topologies with dense neuron connections, the reduction in *N* doesn’t impact the patterns as expected due to sufficient current flow. Finally, the maximum network size used in the current experiment was *N*=10000, because of computational restrictions. However, the current observations of similar patterns in (*α, β*) should hold for larger neuron networks as well. From these comprehensive experiments, we observe that by making the networks larger and with proper upscaling of the stimulating currents, the spiking activity of each neuron does not diminish. Therefore, the fractality induced by network topology in the spiking data can be captured.

### Stability of memory and scale parameters

The neuron particles model also enables tracking functional changes in the network topology across species, tasks, and task epochs. Fig. 4c shows the inferred model parameters (*α, β*) for different motion directions and epochs of the direction discrimination task in the monkey pre-arcuate gyrus. The memory parameter *β* is highly stable (Fig. 4e), compatible with the stability of intrinsic connectivity and the underlying dynamics of neural responses across tasks and task epochs in this region of the monkey cortex (*41*). Since *β* reflects the collective effect of the multiple closed-loops in the topology, we predicted little change across task epochs. The scale parameter *α* which reflects path lengths and in-degree is also stable (Fig. 4d). The stability of *α* and *β* despite obvious changes in the sensory inputs and motor outputs involved in the three distinct epochs suggests the neuron population-level response statistics and network topology are stable. This stability is reminiscent of the random network topology which has been hypothesized to be present in the pre-frontal cortex (*42*) to support adaptive learning and behavior. In contrast, if changes in input and output across epochs were associated with emergence of new loops or changes in path lengths of network (e.g., activation of a hub neuron in a BA network), such stability would be unexpected.

## Discussion

We developed a novel statistical physics inspired model, which captures the causal memory and fractal patterns of inter-spike intervals recorded *in vivo* from single neurons. More precisely, we define the “neuron particle” as the time interval between consecutive spikes, that through its path-based history captures the degree of long-range memory emerging in neuronal activity due to intrinsic local computation and interaction with other networked neurons. We first show how the hierarchical information encoded in the spike trains is captured using the proposed neuron particles model (Fig. 1). Next, we show how the neuron particles model can encode information about the underlying neural networks that were not directly and completely observed (Fig. 3).

Although it is widely recognized that the pattern of spike trains in one specific neuron is a reflection of a network of interacting neurons and glia connected to it, to the best of our knowledge, no previous study has shown that it is possible to predict network topology based on single unit activity. Previous studies largely focused on pairwise correlations or higher order statistical features based on neural population responses (*1, 38*). Although knowledge of network connectivity enables predictions of neural responses features, the converse has been elusive until now (*43, 44*).

The proposed method of neuron particles generalizes to any complex network with interactions between nodes with discrete time intervals between activities. As an example, first, the online hate events which have been linked with several extremist activities (*2*) could be used to infer the underlying terrorist-group network structure. Monitoring the complete network is difficult as such individuals are very discreet. Using the ‘social particles’ analogy, using very few users’ data, the network topology could be inferred using computed fractality of events. Armed with the topology, it may be possible to predict the timing and implicated parties involved in a terrorist attack by detecting activity in only a few of the nodes, and not necessarily the actual perpetrators. As another example, climate change events impact rainforests across the globe. The detailed forest maps (*4*) and tree species-based information could be used to make a time-varying complex network. Extracting the fractality in the climate change events by ‘climate particle’ jumps and observing its variations by changing tree population network, the next damaging event, especially for vulnerable tree species could be foretold. Third, the impact of the introduction of species by humans to the network of interactions among other species brings ecological and evolutionary consequences (*3*). Capturing the change in the network over a global scale is challenging, but using the particles approach, the change in the change in *α* and *β* obtained using a ‘species particles’ jumps from only a handful of nodes can provide a change in the underlying network topology. The time fractality computed over multiscale interactions provides a way to capture human interventions over several years.

Our advance has several important implications for neuroscience applications. The first is that the neuron particles model can be used to analyze single unit data to capture the causal information and long-range memory contained in the statistically self-similar (repeating) patterns of time intervals between spikes for superior inference. The underlying networks of the brain could be inevitably more complex and may not be a direct match with any of the considered network topologies here. But, since the considered networks in Fig. 3d span a wide spectrum of complex networks, they can be treated as canonical networks. Future studies are needed to consider scenarios where the network is a blend of various canonical structures, and whether the neuron particles can extract proportions of these constituent networks.

Second, if features of the underlying neuronal networks can be decoded then theoretically better predictions of behavior should be possible from single unit recordings. Indeed, the real-world examples show how using the neuron particles produces superior predictions of behavior when compared to conventional approaches which use the mean firing rates (Fig. 4). By the same logic, better predictions of neuronal activities should also be possible. If the network can be inferred, then that should constrain possible paths of information in response to new stimuli, or specific neuronal activation patterns. To enable predictions of neuronal responses to new stimuli, future work should develop causal inference techniques to identify composable sets of multi-fractional PDEs for multi-dimensional trajectories. Doing so would enable to study the interactions of neurons explicitly using population spike trains. The working hypothesis of the current paper is that the reflection of interaction with the complete network is present in the spike train of each neuron, which we capture through *α, β*. However, with multi-dimensional modeling, a better prediction could be made of joint neuronal activities, for example, inter-neuron spike correlations.

Third, the neuron particles model can allow insights into network structures that enable how the brain processes information. In the brain, interactions of large neural populations implement computations, decisions, and perceptions. The brain, therefore, does not need to rely on the neuron particle concept for its computations. In fact, it is unlikely that the brain extracts information in an analogous way as our statistical model. However, the neuron particles model provides a way to obtain information about the underlying network structure and dynamics, and in this manner, has the potential to provide new insights into mechanisms of brain function that were not possible before. For example, it may be possible to infer that a group of neurons or glia are disrupted by detecting the change in the network dynamics through a single unit, even if the single unit is intact. Although this speculation would need to be confirmed using experimental data from single units where specific known defects are introduced in a circuit to see how well the neuron particles perform in identifying the defect (i.e., needle in the haystack), the implications are tantalizing. For example, it may be possible to measure a reasonable number of neurons and be able to make predictions about the early onset of neurodegenerative disease, epilepsy, brain cancer or other neurological disorders by detecting a change in the network dynamics without directly measuring the implicated tissues.

## Materials and Methods

### Neuron Particles

A neuron particle trajectory is computed by identifying the repeated patterns in the spike train data at various scales. The spike train data recorded from a single unit is represented as a sequence of inter-arriving intervals T = *τ*_1_*, τ*_2_, …*, τ*_*N*_ which is resulted from N+1 spike times. A particle is termed as unique inter-spiking interval, and for the computational purpose is used as a continuous range of inter-spiking intervals. For complete details of neuron particles process is discussed inthe next section. For a particle *τ*_*p*_ with continuous range 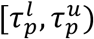, we first define the particle arrival times as

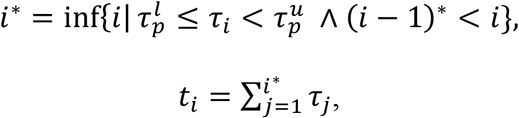

where, *t*_*i*_ is the ith arrival time of the particle *τ*_*p*_ in the inter-arrival time sequence T and 0^∗^ = 0. Next, using the arrival times, we now define the neuron particle trajectory as follows.

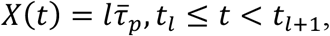

where, 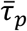 can be defined as average particle size with 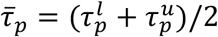 and the start time is taken as *t*_0_ = 0. The multiple trajectories are used as input for the FPDE estimation Algorithm (*31*) using moments approach.

### Neuron Particles Process

The neuron particles process (NPP) aims at obtaining a computational version of the neuron particles from the spike trains. For a scenario where the inter-spike intervals (ISIs) belong to a set of finite cardinalities, for example, Cantor spikes in Fig. 1, the neuron particles can be uniquely identified. However, for a general setup, such assumptions do not hold and ISIs are some real numbers, hence, identifying unique ISIs (or neuron particles) is not computationally useful. Therefore, we resort to ‘range of ISI’ as ‘neuron particles’, for the sake of computation, as we discussed in the previous section. We now present a novel algorithm (NPP) that computes neuron particles from the spike train data which is also pictorially shown in the Fig. 2.

### NPP Algorithm

Initialization: Given a spike train series with the sequence of inter-spike intervals (ISI) as *T* = *τ*_1_*, τ*_2_, …*, τ_N_*, a binning algorithm 𝒜, an initial number of neuron particles *n*_*i*_, lower/upper acceptable values of ISI frequency as *lb/ub*.

Step-1: Using algorithm 𝒜 with number of bins *n*_*i*_ get the histogram of ISI sequence *T* with bin boundaries 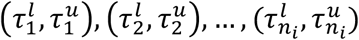 and bin counts 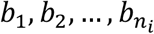. The constraint on data sequence *T* is such that *b*_1_ ≥ *lb*, otherwise terminate the NPP algorithm.

Step-2: For *i* = 1,2, …*, n_i_* if *b*_*i*_ > *ub* then

For the data subsequence 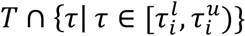 get the histogram with number of bins as 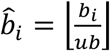.

Update the histogram of the sequence *T* with the new bin edges and bin counts. This is termed as ‘split’ step.

The updated histogram of *T* has *m*_*u*_ ≥ *n*_*i*_ bins with bin counts 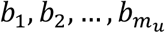.

Step-3: With *i* = *m*_*u*_, while *i* > 1, if *b*_*i*_ < *lb* then

Start with *i*th bin and ‘merge’ consecutive bins such that merged bins have bin counts exceeding *lb* or with 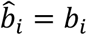 iterate over *j* = *i* − 1*, i* − 2, …,1 as 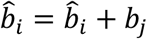. till 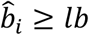.

Update the new bin boundaries as 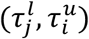 with bin count 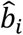 and set *i* = *j* − 1. This is termed as ‘merge’ step.

The resulting histogram after merging has total bins *m* with bin counts *b*_1_*, b*_2_, …*, b_m_*.

Step-4: Output the final iterated bin edges 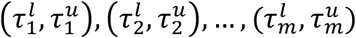. The bin edges are referred as neuron meta-particles which consists of neuron particles such that *i*th meta-particle represents a group of particles with size ranging between 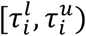.

The initial number of particles *n*_*i*_ is guessed based on the length of the input spike train data *N*. The algorithm iteratively updates the number of particles based on the provided bounds *lb/ub*. While any value of *lb/ub* is feasible if 1 ≤ *lb* ≤ *ub* ≤ *N* ideally, we wish *lb* = 1 and *ub* = *N* such that there is no constraint on the neuron particles. However, we found that setting a low value of lower-bound (for example, *lb* = 1) results in spurious ISI intervals being selected as neuron particles which never repeats in the ISI sequence *T*. Such ISIs could be due to the experimental noise and are not directly useful. We have chosen *lb* = 5 in all the experiments. The choice of *lb* = 5 is made by cross-validating over the simulated data of the Fig. 2 such that ISIs (within a small margin) to be selected as neuron particles repeats in the given input sequence *T*.

### Cantor Spikes

The Cantor spikes concept is used for demonstrating the fractal memory in the spike trains. The Cantor set is constructed by repeatedly deleting 1/3 of the unit interval. The extension of Cantor set, C_r_ is defined such that at any iteration-n, rth ratio is removed from each subsection. The extended Cantor set C_r_ with ratio r at each depth level N has deleted intervals of size r, *rδ, rδ*^2^, …, *rδ^N^* ^−1^, where *δ*=(1-r)/2. For example, at depth N=2, the sequence of deleted intervals is T =*rδ*, r, *rδ*. For the computational purpose, at any depth N, the sequence is generated by constructing a binary tree with root node having weight r, and each child (left and right) is assigned a weight of (1-r)/2 times the weight of its parent. The sequence T is obtained by traversing the binary tree with left child first rule. We note that, for each depth N, the neuron particles can be analytically identified as r, *rδ, rδ*^2^, …, *rδ^N^* ^−1^. The fractality of Cantor sets (and hence cantor spikes) is analytically captured via box counting dimension (*35*). The box counting dimension for set A is defined as

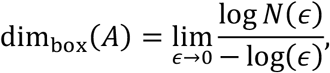

where, *N*(*ϵ*) are the number of boxes with diameter *ϵ* required to cover the set A. At iteration-n, there are 2^n^ resulting sub-intervals, each of size 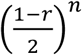, therefore, the box-dimension is written as

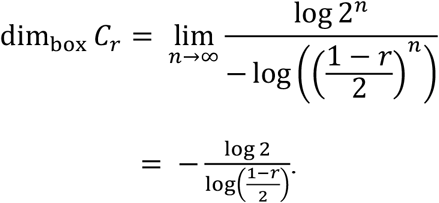

### Artificial Neuronal networks

For analyzing the network influence, the first ingredient is the neuron transfer function which we implement through a non-linear Izhikevich model (*45, 46*) with axonal conduction delays and spike timing dependent plasticity. A total of 1000 neurons were simulated for 60 sec using 80:20 ratio of excitatory and inhibitory neurons (*46*) in Fig 2. In Fig. S3, using a similar setup, the network size is varied from N=1000 to N=10000. The second ingredient of network topologies is described next in this section. For every topology, 50 network realizations were generated and the fractal memory, scale results were concatenated for the purpose of box plots.

#### Random Networks

The Erdös-Rényi (*39*)(ER) model of a random network connects each pair of nodes with association probability *p*. The node-degree is a binomial distribution with parameter *p*.

*Scale-free networks* are the network topology with degree distribution following a power-law, i.e., the tail probability for degree *d* is *P*(*d*) ~ *d*^−*γ*^, where *γ* is the degree exponent. The Bárabasi-Albert (*47*)(BA) model associates a node to other nodes with a probability proportional to their in-degrees, commonly referred to as rich-gets-richer phenomenon. The generated network has *γ* = 3. The Dorogovtzev(*48*) extension of BA model associates a node with *m* edges to other nodes with probability proportional to the sum of attraction parameter *a* and in-degree. The resulting scale-free network has tunable degree exponent with an attraction parameter *γ* = 2+*a*/*m*.

*Multi-fractal networks,* unlike random networks, possess hierarchal, fractal, and self-similar properties. The multi-fractal network (MFN) generator using a base measure P (m×m) and unit interval L (m×1) successively generate deeper measures at depth K which are self-similar (*49, 50*). Compared to BA models which are mono-fractal, the multi-fractality generalizes the notion and mimic real-world networks. In the experiments, we take m=2 and a base measure of *P* = [0.6,0.5;0.5,0.4]. The change in length measure from 0.1 to 0.9 drifts the average degree from low to high due to higher value of *P*_11_ compared with *P*_22_ as we see in Fig.3c.

### Varying-size of the network

The baseline case for the network size is N=1000, as taken in the experiments for Fig. 2. For simulating the smaller/bigger networks following the same topological structures as in Fig. 2, we normalize the in-degree distributions accordingly. The normalization procedure is carried out to avoid build-up of large number of incoming connections as the network size grows/loss of connections when network size decrease. Next, to maintain the same proportion of stimulating currents in the network as in the baseline case, we also normalize the current distribution across each 1sec (for a total of 60sec). The currents are bombarded in the ratio of the network bigger/smaller than the baseline case, with high currents in the bigger networks and shallow currents in smaller networks.

### Node-based Multi-fractal spectrum

The fractal structures in a complex network can be quantified using multi-fractal spectrum (MFS) (Fig. S4a) analysis. A common method to perform such analysis is by using the box-covering approach. The given network (possibly weighted) is fractal if the box-size and the number of nodes covered by the box in the given graph obeys the relationship: 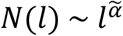, where 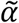 is Lipschitz-Holder exponent, *N(l*) is the number of nodes covered by the box of size *l*. The exponent 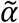 represents the mono-fractal dimension of the structure. Instead of mono-fractal, the given network might have multiple scaling structures which are present together. The given network is multi-fractal if the box count relationship satisfies: 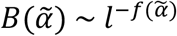, where 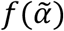 is the singularity or multi-fractal spectrum, 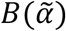 is the number of boxes with exponent 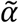. From the computational viewpoint, a distortion exponent *q*(*q* ∈ ℝ) is used to weigh different exponents 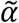. For example, negative *q* endorses exponents that have smaller box-counts while positive *q* for boxes with larger number of nodes. The distortion exponent and mass exponent *τ*(*q*) together are used to write the fractality relationship as follows: ∑_*i*_*N*_*i*_ (*l*)^*q*^ ~ *l*^*τ*(*q*)^, where *N*_*i*_ (*l*) is the number of nodes in the *i*th box of size *l*. The pair (*τ*(*q*)*, q*) is related to 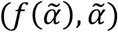 using the Legendre transformation as follows.

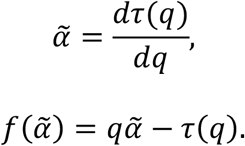

For the neuron networks in Fig. 3, to get the multi-fractal spectrum we have used a node version of the multi-fractal spectrum (*51*). For the box-counting, the distance between a pair (*i, j*) of neurons is taken as 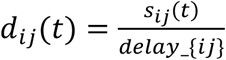, where *s*_*ij*_(*t*)is the time-varying conduction strength of the neuron link, *delay*_*ij*_ is the conduction delay. This combines the effects of both conduction strength and delays on the current flow across synapses. The MFS analysis in Fig. S4b-d is presented over complete simulation period of 60 sec.

## Author contributions

G.G. conceived the neuron particles concept and conducted all the experiments. G.G., J.S., and P.B. designed the simulation experiments. R.K. collected the monkey data. G.G, R.K., and P.B. designed the monkey data experiments. All authors discussed the results and implications, and contributed extensively to the multiple subsequent revisions of the paper.

## Competing interests

The authors declare no competing interests.

## Data availability

The data generation scheme for the simulation studies will be available publicly upon publication. The Monkey dataset is publicly available at http://www.cns.nyu.edu/kianilab/Datasets.html.

## Supplementary Materials

**Fig. S1.**
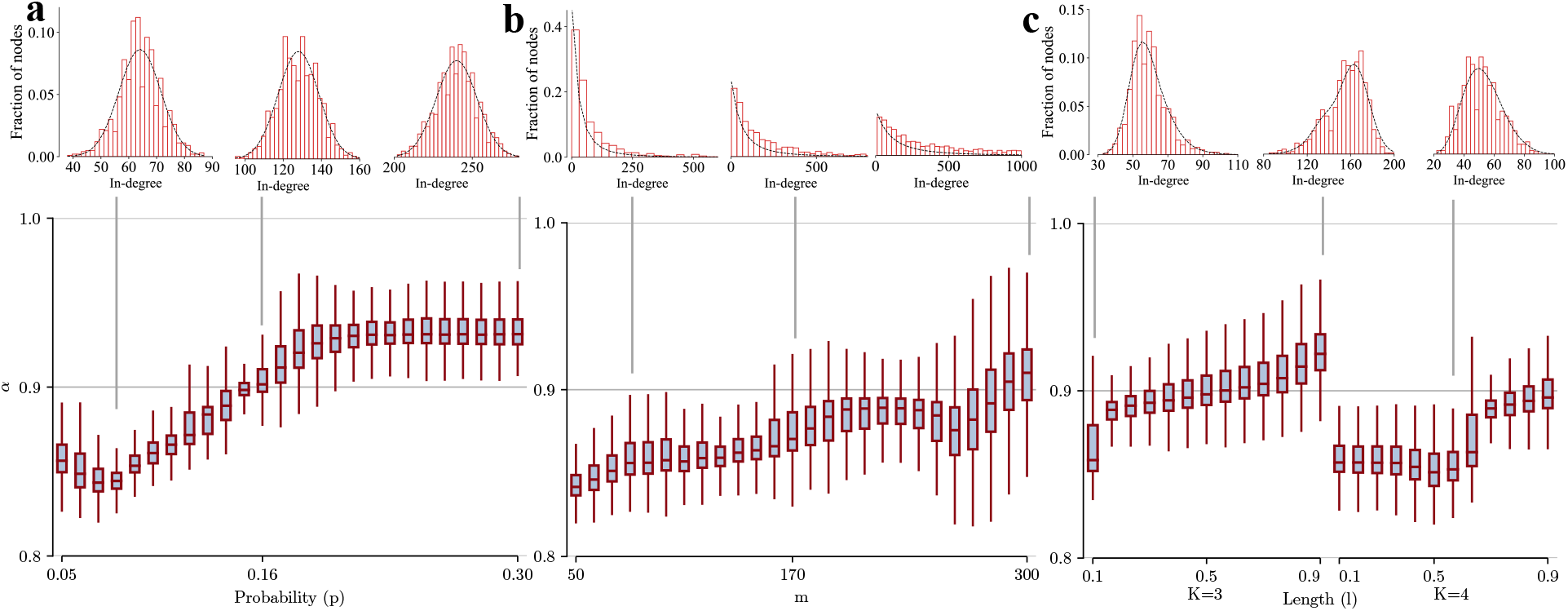
Network topology influence on scale. a, Random network with probability of existence of edge being *p*. The probability varies linearly from 0.05 to 0.30 and consequently increases the average node-degree. The excitatory neuron node-degree scaled histogram is in red bars and the black curve fits the histogram with Gaussian distribution. The neuron population scale parameter *α* variation with p is represented as box plots. The line across the box plot indicates the median, and upper and lower hinges indicate 75^th^ and 25^th^ percentiles, respectively. The whiskers extend to the lowest and the largest values but within 1.5xIQR. b, Dorogovtsev (*46*) extension of Bárabasi-Albert (*47*) scale-free Network with outgoing links *m* and attraction parameter *a* with *a=m*. The excitatory neuron node-degree scaled histogram is in red bars and the black curve fits the histogram using analytical expression (*48*). The outgoing links *m* varies from 50 to 300, and the variation of population scale parameter *α* is shown as box plots. c, Multi-fractal network (*49, 50*) with base measure P = [0.6,0.5;0.5,0.4] and length measure *l* variation linearly from 0.1 to 0.9 for two values of depth K = 3, 4. The excitatory neuron node-degree scaled histogram is in red bars and the black curve fits the histogram with the analytical expression of degree distribution (*49*)(*49*). The neuron population scale parameter *α* is represented as box plots.

**Fig. S2.**
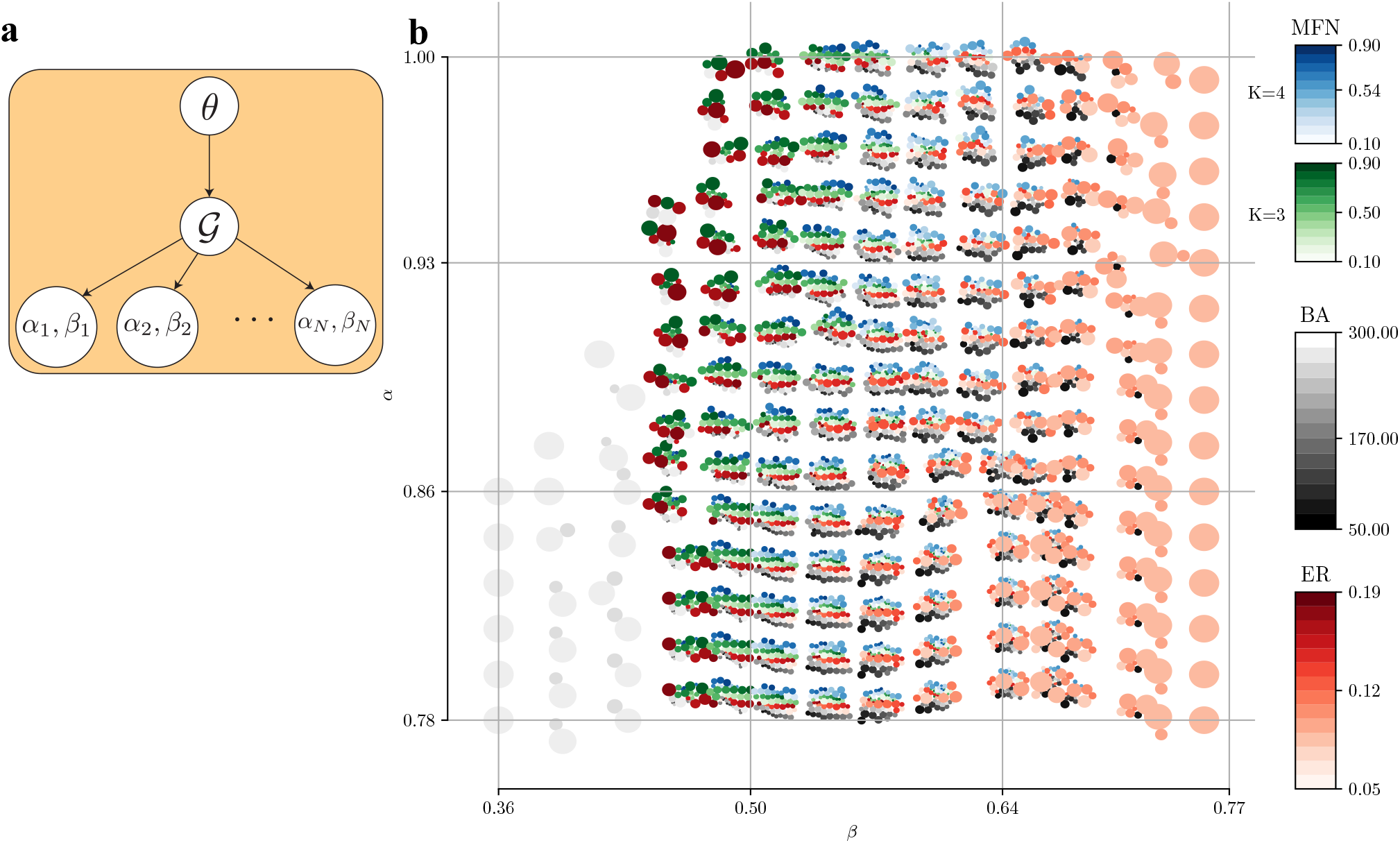
a, Bayesian graph model for the neuronal network (with N neurons) and extracted parameters (*α,β*) for each of N neurons. b, Prediction spectrum of the networks using *α* and *β* computed from a single spike train for synthetic neuron networks as discussed in Fig.3 in the main text. For each tuple (*α,β*) the confidence that it belongs to a particular network type is indicated by its corresponding bubble size.

**Fig. S3.**
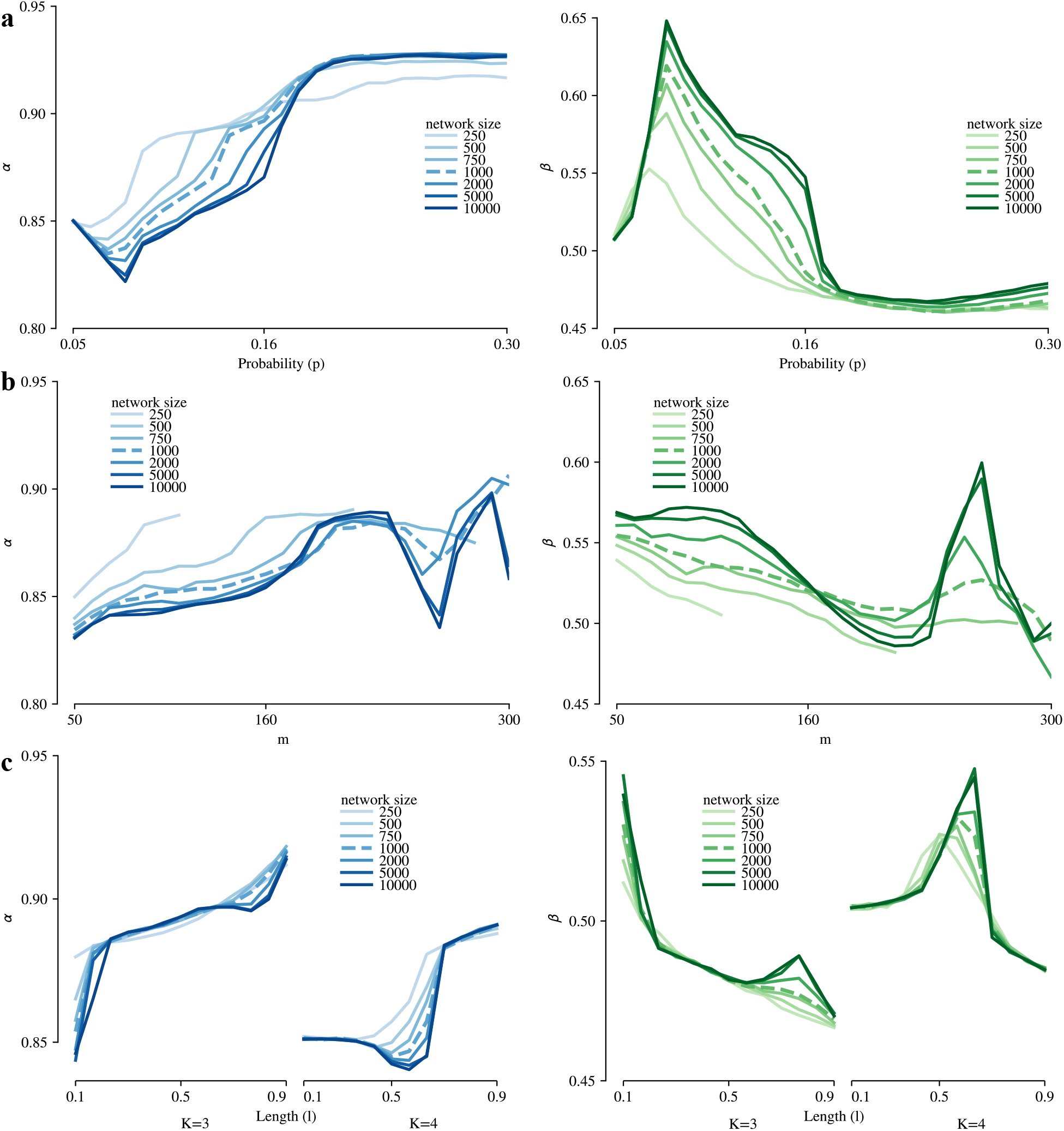
Memory and scale invariance to the network size. a, Erdos-Renyi random network with probability of existence of edge being *p*. The probability varies linearly from 0.05 to 0.30. b, Dorogovtsev (*46*) extension of Bárabasi-Albert (*47*) scale-free Network with number of outgoing links *m.* The outgoing links number *m* varies from 50 to 300. For the smaller networks N=250, 500 and 750, the larger values of *m* are not feasible to simulate a power-law degree distribution. The highest value of m is limited as m=110, 210, 290 for N=250, 500, 750, respectively. c, Multi-fractal network (*49, 50*) with base measure P = [0.6,0.5;0.5,0.4] and length measure *l* variation linearly from 0.1 to 0.9 for two values of depth K = 3, 4. The median of node parameters (scale *α*, memory *β*) is shown for different network size for each network topology in a, b, c.

**Fig. S4.**
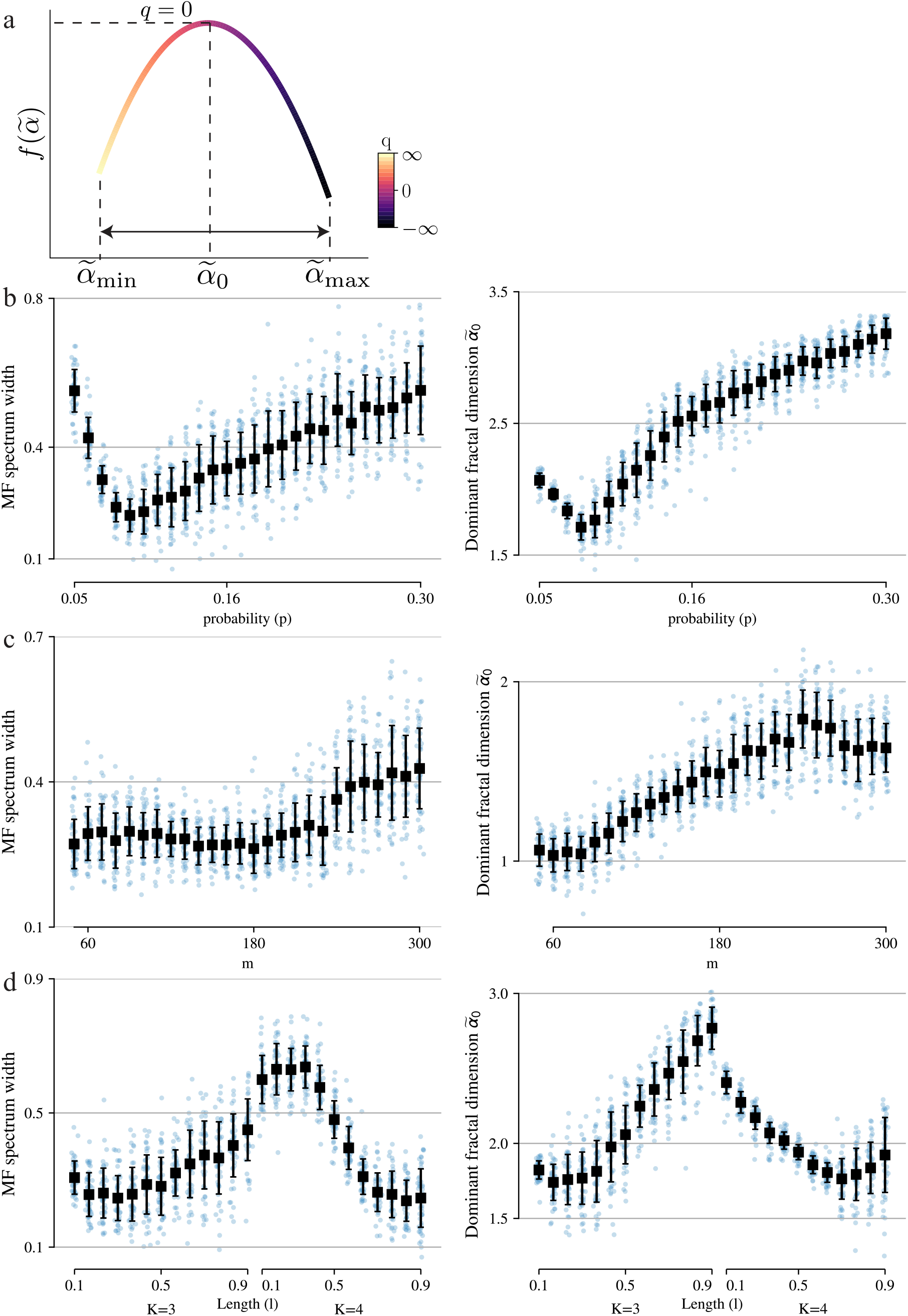
a, Multifractal spectrum for a weighted directed graph. The dominant fractal dimension is 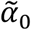and the spectrum width is 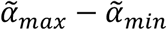. b, Multifractal spectrum width and dominant fractal dimension for Erdos-Renyi networks parameterized with connection probability (p). The probability varies linearly from 0.05 to 0.30 as in Fig. 3a. The black square represents mean and vertical lines represent s.d. for the corresponding network parameter. c, d, Same plots for Barabasi-Albert networks in c and Multi-fractal networks in d. The network parameter varies as in Fig.3b,c, respectively.

**Fig. S5.**
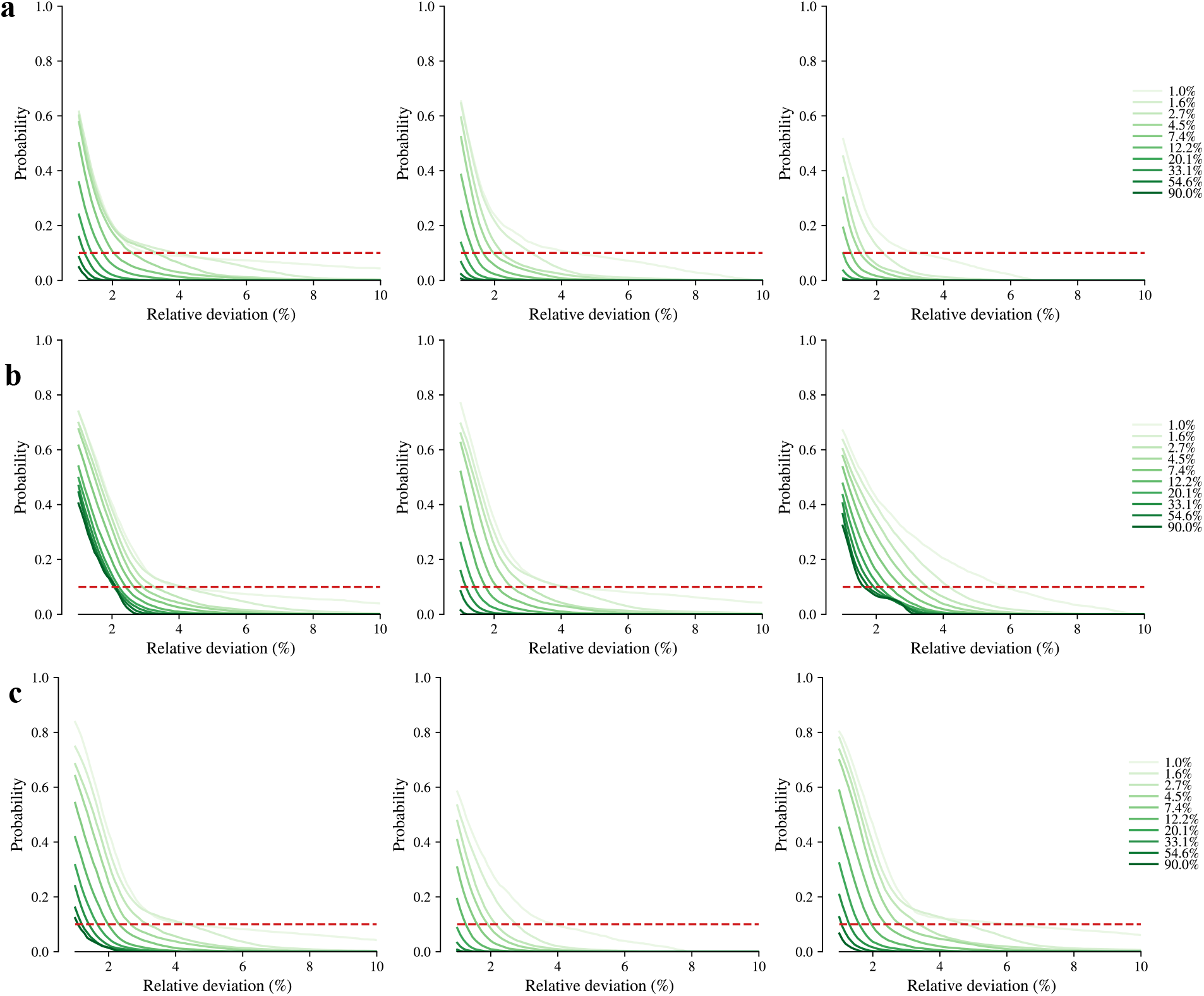
a, Probability of relative deviation of a partial sample (*δ*%) of *α* from the neuron population of Erdos-Renyi networks with p=0.08, 0.16, 0.3 from left to right. The red line is 90% marker for error such that curve values below red line denotes >90% confidence in the relative deviation. b, Same for scale-free Barabasi-Albert networks with m=80, 170, 300 from left to right. c, For Multi-fractal networks with (K,l)=(3,0.1), (3,0.9), (4,0.6) from left to right.

**Fig. S6.**
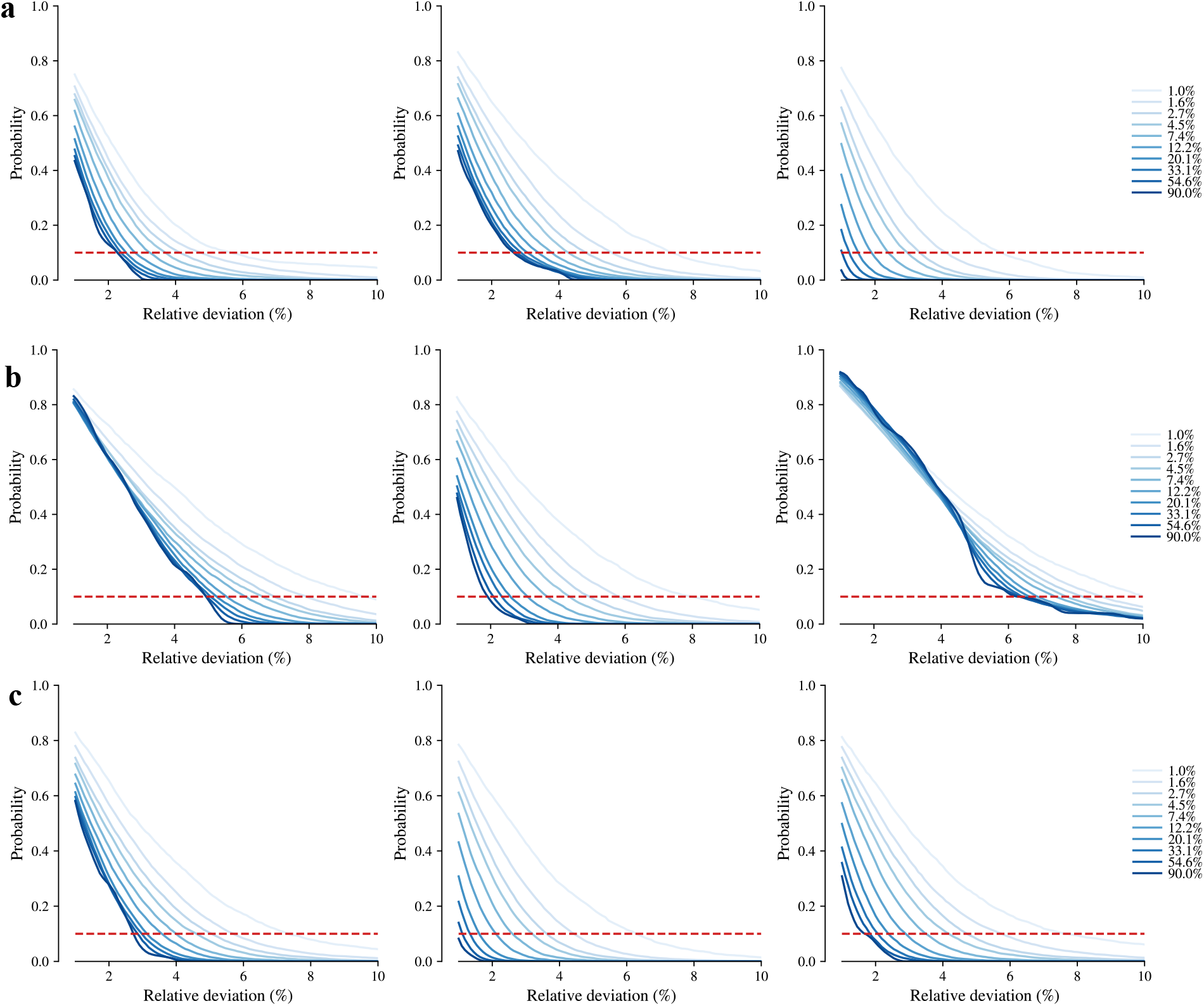
a, Probability of relative deviation of a partial sample (*δ*%) of *β* from the neuron population of Erdos-Renyi networks with p=0.08, 0.16, 0.3 from left to right. The red line is 90% marker for error such that a curve with corresponding *δ* below red line denotes >90% confidence in the relative deviation. b, Same for scale-free Barabasi-Albert networks with m=80, 170, 300 from left to right. c, For Multi-fractal networks with (K,l)=(3,0.1), (3,0.9), (4,0.6) from left to right.

**Fig. S7.**
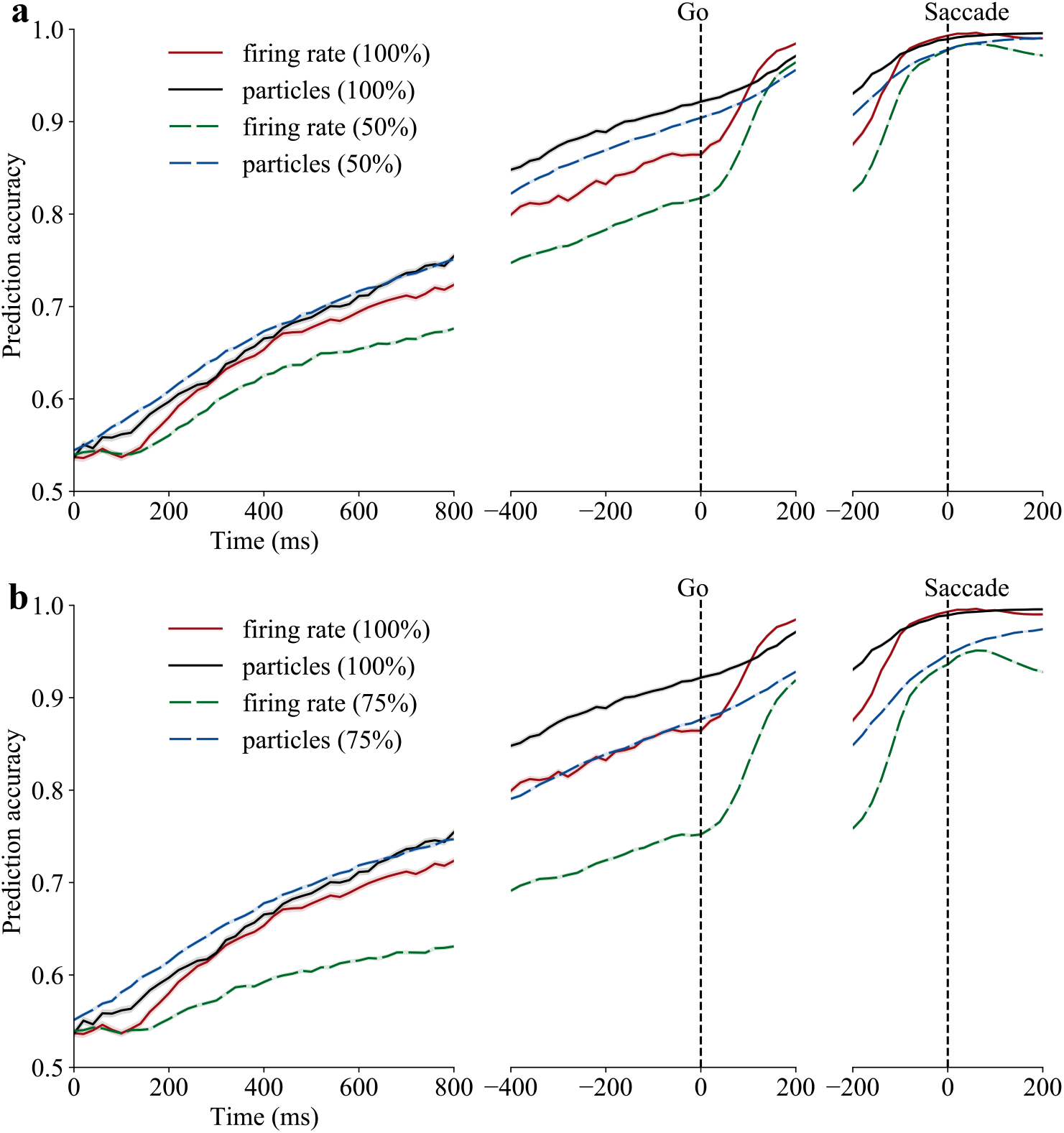
Neuron particles with partial spiking data. a, Prediction accuracy gap between neuron particles and mean firing rate approach increases under partial data. The partial data setup is mimicked by uniformly random sampling of 50% of the spikes from each single unit. The random experiment is repeated 5 times and the results are averaged. Average prediction accuracy mean ± s.e.m. (across sessions and resampling) over time with classifier setup same as Fig. 4. b, same as a but selecting 25% spikes from each single unit uniformly at random.

**Fig. S8.**
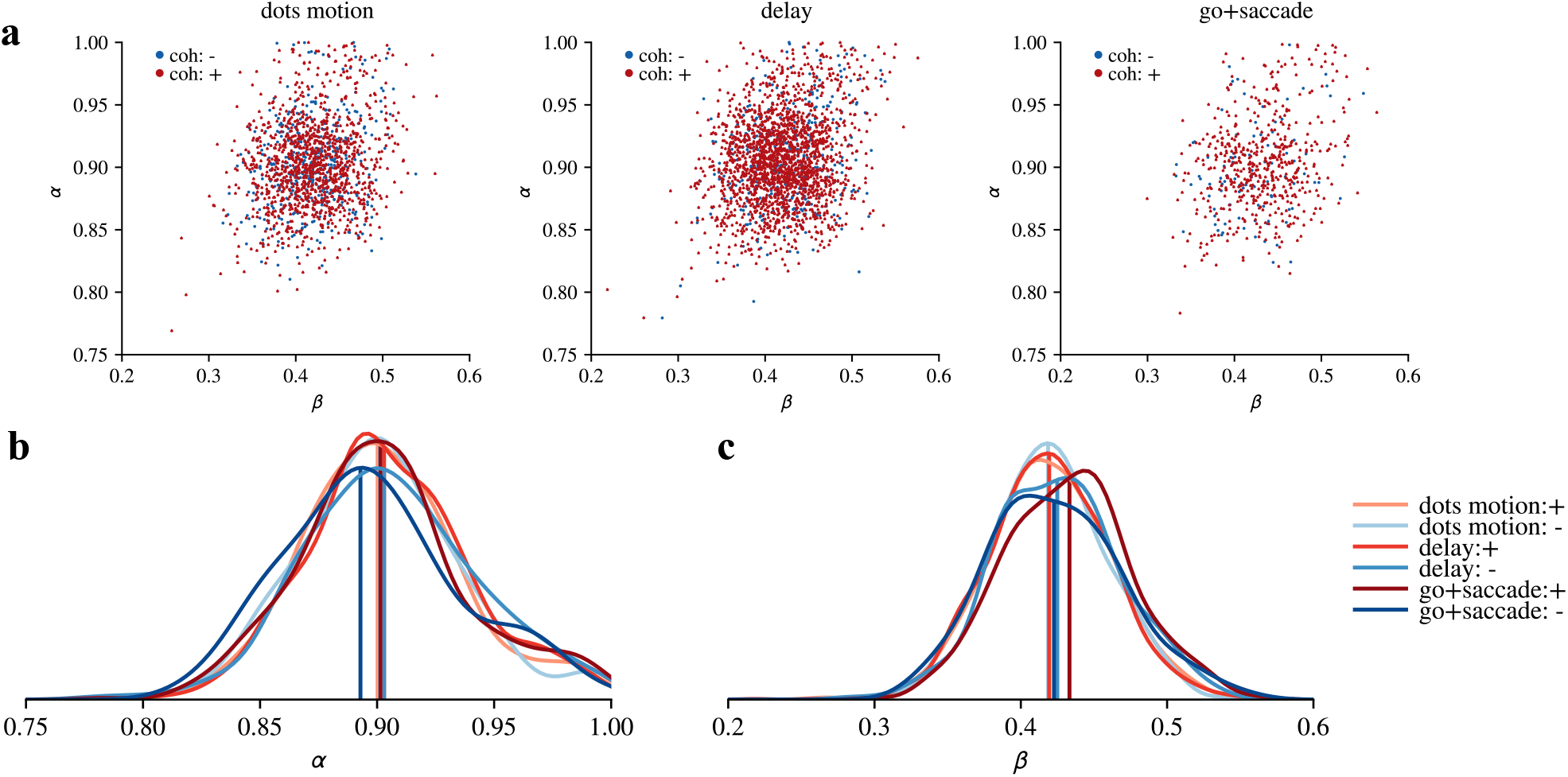
a, The estimated parameters for monkey 2 across three epochs: motion viewing (200-800 ms after motion onset), delay (200 ms after dots offset to −50 ms from go cue), and go+saccade (go cue to 200 ms after saccade initiation). The parameter tuple was separately calculated for motion in favor and against the neuron’s preferred saccade (+ and – motion coherence, respectively). b, c Gaussian-kernel smoothed densities for *α, β* across three epochs and 2 coherence levels with straight line being the median.

## Notes

### Competing Interest Statement

The authors have declared no competing interest.

